# Uncertainty-Guided Decision-Making in Bumble Bees

**DOI:** 10.64898/2026.09.15.751944

**Authors:** Yingying He, Qiao Ye, Lin Lin, Runzhu Yuan, Qian Wang, Song Wu, Song Chen, Lin Yuan

## Abstract

The evolutionary origins of cognitive monitoring remain contested. Metacognition, the capacity to monitor one’s own cognitive states, has long been linked to complex vertebrate brains, yet whether a miniature brain can evaluate the reliability of its internal representations is unknown. Here we demonstrate that bumble bees engage in uncertainty-guided decision-making. Bees dynamically adjusted opt-out choices according to perceptual difficulty, settling for a smaller guaranteed reward to avoid errors, and actively paid a reward cost to seek information under uncertainty. Without any retraining, bees transferred this ‘opt-out-under-uncertainty’ rule to novel tactile and working-memory tasks under an all-probe design. This strategy was stable across individuals and accurately captured by a confidence-based decision model. Our findings suggest that a brain of approximately one million neurons can support an abstract domain-general uncertainty-monitoring policy, indicating that the neural substrates for uncertainty-guided decision-making may be far more ancient and widely distributed than previously assumed.

## INTRODUCTION

The cognitive revolution in invertebrate research has steadily dismantled the view that sophisticated cognition requires a large vertebrate brain (*1*). Social insects, once regarded as reflexive automatons, are now recognized as flexible learners whose cognitive capacities rival those of some vertebrates (*2, 3*). Bumble bees (*Bombus terrestris*) have been central to this shift, with recent studies showing that they can exhibit positive affective contagion, spontaneous problem-solving, and learn abstract rhythmic structure that generalizes across several sensory modalities (*4–6*). Complementary work on honey bees has further revealed that they socially transmit foraging information through the waggle dance (*7*). Collectively, these findings demonstrate that brains with fewer than one million neurons can support rich internal representations, social sensitivity, cross-modal abstraction, and even symbolic communication.

Yet this revolution has largely concerned cognitive content—what an insect brain can represent, compute, and transfer. A deeper, more fundamental question concerns cognitive monitoring: whether an insect can evaluate the reliability of its own representations. Metacognition, broadly defined as the ability to monitor and regulate one’s own cognitive states, is considered by some theories to be a core computational component of consciousness, self-evaluation, and higher-order thought (*8–10*). Behavioral evidence consistent with metacognition has been reported primarily in vertebrates with highly encephalized brains, including primates, dolphins, and corvids (*10–14*). This distribution has supported the assumption that the neural architecture required for internal uncertainty monitoring emerged within the vertebrate lineage and depends on substantial cerebral complexity. Evidence for metacognition in an invertebrate would therefore challenge this big-brain account of the biological prerequisites for cognitive monitoring (*15–17*).

Bumble bees provide an ideal test case because their known cognitive capacities already extend beyond simple associative learning. If bees can form abstract and amodal representations, they may also possess mechanisms for evaluating when those representations are unreliable (*13, 15*). The key challenge is to distinguish such monitoring from simpler explanations, including task difficulty, stimulus avoidance, reward history, and motor habit. For instance, previous work showed that bees avoided difficult choices, but the behavior could be explained by stimulus-specific avoidance learning (*18*). We aimed not only to test for behavior consistent with uncertainty monitoring but to interrogate its nature: is it a domain-specific learned association, or an abstract cognitive rule deployable across novel contexts?

To address this, we designed an all-probe behavioral framework that operationalizes uncertainty monitoring in freely foraging bees. In a visual discrimination task of graded difficulty, bees were given a persistent safe exit that yielded a smaller but guaranteed reward. This allowed uncertainty to be expressed as an economically meaningful choice: commit to the discrimination when confidence is high, or accept lower-value certainty when confidence is low. To test whether bees’s behavior is consistent with an active assessment of insufficient evidence, we further asked whether they would actively seek information at a reward cost. Finally, to distinguish an abstract uncertainty-monitoring rule from a task-specific heuristic, we tested whether bees could transfer an opt-out-when-uncertain strategy to novel tactile and memory-based decisions without any retraining.

This approach fundamentally shifts the question from whether insects can represent the world to whether they can monitor the fidelity of those representations. By integrating opt-out behavior, costly information seeking, and cross-domain transfer within a single all-probe design, this study tests whether a miniature brain can implement a core computational capacity for uncertainty monitoring with far fewer neurons than previously assumed (*19–21*).

## RESULTS

### Bumble bees exhibit an opt-out response under perceptual uncertainty

We developed an all-probe behavioral paradigm to decisively test whether bumble bees use uncertainty to guide choice. Individual foragers (*N* = 192) were trained in a flight tunnel to discriminate between two vertical gratings that differed in spatial frequency. Correct choices yielded 30% sucrose, whereas errors were punished with 1 mM quinine. Easy (e.g., 0.1 vs. 0.3 cycles/mm) and Hard (e.g., 0.1 vs. 0.12 cycles/mm) trials were randomly interleaved, creating a graded perceptual difficulty structure. After performance stabilized above 80% correct on Easy trials (Fig. S1 and Table S1), bees entered the test phase.

In this phase, we added a third, permanently available gray platform that functioned as a safe exit. During brief shaping (≤10 trials per bee), punishment was suspended and landing on the gray platform rewarded with 15% sucrose, teaching bees that the safe exit offered a reliable but lower-value option. In the formal test, full reward and punishment contingencies were reinstated for the discrimination task (Fig. 1A), forcing a three-choice decision: discriminate or exit.

**Fig. 1.**
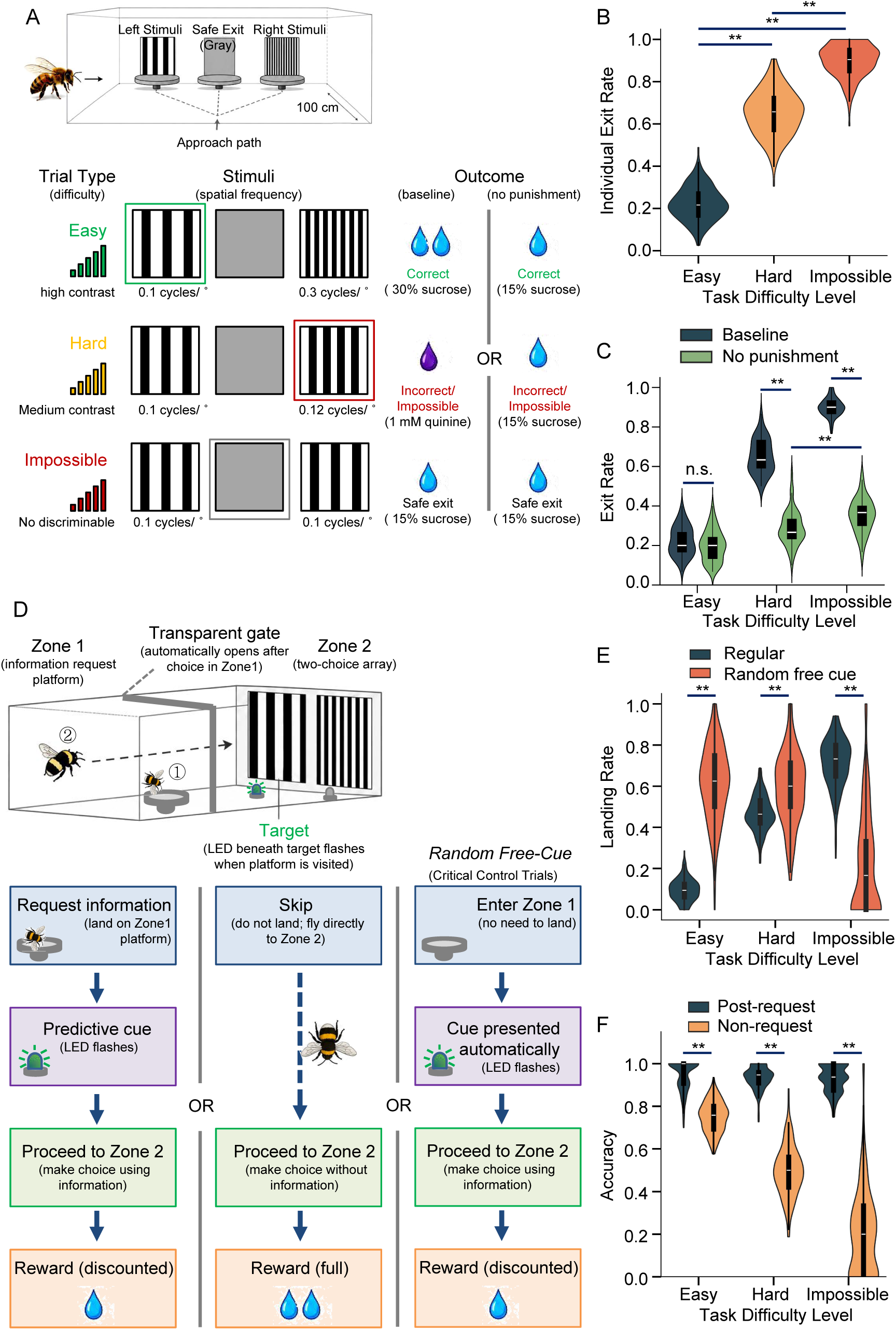
Bumble bees show difficulty-dependent opt-out and actively seek information under uncertainty. (A) Experimental paradigm. Schematic of the linear flight tunnel. Bees initiated trials from a start gate and flew to a decision zone with three landing platforms: two outer platforms displaying vertical gratings (varying spatial frequency) and a central permanent “safe exit” (uniform gray disk). Correct choices were rewarded with 30% sucrose; errors punished with 1 mM quinine; the safe exit yielded a smaller but guaranteed 15% sucrose reward. Trial difficulty was titrated: Easy (high discriminability, e.g., 0.1 vs. 0.3 cycles/mm), Hard (low discriminability, e.g., 0.1 vs. 0.12 cycles/mm), and Impossible (identical 0.1 cycles/mm gratings). (B) Opt-out rates calibrated to task difficulty. Bees opted out more on Hard (65.1 ± 10.7%) and Impossible trials (89.9 ± 7.7%) than on Easy trials (22.3 ± 7.9%) (GLMM, Hard vs. Easy: *Z* = 46.2, *p* < 0.001; Impossible vs. Hard: *Z* = 25.6, *p* < 0.001). (C) Punishment removal control. A separate cohort (*n* = 60) completed the same paradigm under baseline (with punishment) and no-punishment removal (all outcomes: 5 µL 15% sucrose) conditions. Removing punishment nearly abolished the difficulty gradient (Hard vs. Easy: *p* = 0.09), while Impossible trials still elicited modestly higher opt-out than Hard trials (*p* < 0.001). **, *p* < 0.001; n.s., not significant. (D) Two-stage information-seeking paradigm. In Zone 1, bees could land on an “Information Request” platform, which triggered a 500-ms flashing cue indicating the correct stimulus in Zone 2. Requesting information incurred a reward cost (15% vs. standard 30% sucrose). (E) Information requests scaled with uncertainty. Landing rates: Easy (9.2 ± 3.0%), Hard (45.6 ± 6.8%), Impossible (70.8 ± 10.1%) (GLMM, Hard vs. Easy: *Z* = 37.1, *p* < 0.001; Impossible vs. Hard: *Z* = 22.6, *p* < 0.001). (F) Information seeking resolved uncertainty. After requesting information, bees achieved uniformly high discrimination accuracy (>92%) across all difficulty levels.

Bees chose the safe exit in a difficulty-dependent manner: opt-out rates were 65.1 ± 10.7% on Hard trials versus 22.3 ± 7.9% on Easy trials (mean ± SD; generalized linear mixed model, GLMM, *Z* = 46.2, *p* < 0.001; Fig. 1B and Fig. S2A). When bees chose to discriminate on Hard trials, accuracy remained significantly above chance level (71.8 ± 11.6%; binomial test, *p* < 0.001; Fig. S2B and Table S2). This pattern indicates that bees accepted risk when the available evidence was sufficient to support a choice, while selecting the safe exit when uncertainty was higher.

To further characterize the nature of this behavior, we examined additional behavioral correlates. Bees exhibited longer decision latencies and more antennal investigations on Hard than Easy trials (Fig. S2C-F, and Table S3), consistent with increased evidence gathering under uncertainty. Moreover, in an Impossible control condition with identical gratings, opt-out rates reached 89.9 ± 7.7%, significantly higher than on Hard trials (GLMM, *Z* = 25.6, *p* < 0.001; Fig. 1B), and 187 of the 192 bees showed higher opt-out on Impossible trials than on Hard trials (binomial test, *p* < 0.001; Table S2). This response pattern demonstrates that bees scaled their opt-out behavior to the level of uncertainty they experienced.

In a separate forced-choice control experiment without the safe exit (*N* = 60), bees exhibited a small absolute difference in decision latencies between Impossible and Hard trials (2604 ± 356 ms vs. 2506 ± 302 ms), chance-level accuracy on Impossible trials (50.8 ± 4.7%; binomial test, *p* = 0.42), and uniformly low mid-flight return rates across all difficulties (all < 3%; Table S4). These findings collectively rule out a pure response - conflict account and instead support an uncertainty-guided decision process.

To further distinguish uncertainty monitoring from stimulus-driven avoidance, we eliminated punishment in a separate cohort of visually trained bees (*N* = 60) while keeping all outcomes equally rewarded (5 µL 15% sucrose; Fig. 1A). Under this no-punishment condition, opt - out rates on Hard and Impossible trials dropped sharply (both *p* < 0.001), and the Easy-Hard gradient was substantially attenuated (28.9 ± 10.1% vs. 19.9 ± 9.3%; GLMM, *Z* = 1.67, *p* = 0.09; Fig. 1C). However, Impossible trials still elicited higher opt-out rates than Hard trials (35.1 ± 13.4% vs. 28.9 ± 10.1%; GLMM, *Z* = 5.02, *p* < 0.001; Fig. 1C and Table S5). When bees committed to a discrimination on Hard trials, accuracy remained above chance (61.5 ± 12.4%; binomial test, *p* < 0.001), whereas accuracy on Impossible trials was near chance (50.3 ± 3.1%; *p* = 0.78; Fig. S2G and Table S5). These results provide strong evidence that bees’ opt-out behavior is driven by uncertainty monitoring rather than stimulus aversion or reward-based revaluation.

### Bumble bees pay a cost to reduce uncertainty

To test whether bees actively recognize when additional information is needed, we designed a two-stage information-seeking task (Fig. 1D). The flight tunnel contained two sequential zones: in the first, a single Information Request platform was available; landing on it delivered no reward but triggered a 500-ms cue (a 5-Hz flash beneath the correct target) in the second zone, making the upcoming correct choice salient. The bee then proceeded to the second zone for a mandatory two-choice discrimination. Correct choices after an information request were rewarded with 15% sucrose, imposing an explicit reward cost relative to the standard 30% for correct choices without a request (Fig. 1D). Thus, the platform had no direct reward value—its sole function was to reveal the correct answer before a required decision.

Bees robustly scaled their information-seeking actions to uncertainty levels, with landing rates of 9.2 ± 3.0% on Easy trials, 45.6 ± 6.8% on Hard trials, and 70.8 ± 10.1% on Impossible trials (GLMM, Hard vs. Easy: *Z* = 37.1, *p* < 0.001; Impossible vs. Hard: *Z* = 22.6, *p* < 0.001; Fig. 1E and Fig. S3). When they did request information, bees used the cue effectively, achieving uniformly high subsequent discrimination accuracy across all difficulty levels (all > 92.5%; Fig. 1F and Table S6).

A free-cue control condition, in which the cue was presented automatically and independently of the bee’s landing decision (Fig. 1D), provided a critical dissociation. Under this condition, bees maintained high accuracy across all difficulties (Fig. 1F), but their landing rates on Easy and Hard trials increased significantly relative to the cost condition (Fig. 1E). In stark contrast, landing rates on Impossible trials dropped markedly (Fig. 1E). This striking dissociation reveals that bees sought information strategically, valuing it only when useful—a pattern opposite to what would be expected if they were merely attracted to the cue itself.

Together, these results show that bees invested effort and accepted a reduced reward to obtain information when discrimination uncertainty was high, consistent with an active uncertainty monitoring rather than passive avoidance or habitual cue-seeking.

### Bumble bees transfer the opt-out strategy across sensory and cognitive domains

To test whether bees had acquired a domain-general strategy for managing uncertainty, rather than a task-specific association, we asked whether the opt-out-when-uncertainty rule could transfer to novel sensory and cognitive domains (*22*).

We first examined cross-modal transfer to touch. A subset of visually trained bees (*N* = 60) was transferred to a tactile discrimination task in which visual gratings were replaced by two vibrating platforms (Fig. 2A). Tactile difficulty paralleled the visual structure: Easy (10 Hz sine wave vs. broadband noise), Hard (sine wave vs. frequency-jittered sine wave), and Impossible (two identical jittered waves). Bees received no tactile training, and all choices—including the safe exit—yielded the same small reward throughout the test session. This all-probe design prevented any new associative learning, ensuring that structured opt-out behavior could reflect a strategy carried over from the visual task.

**Fig. 2.**
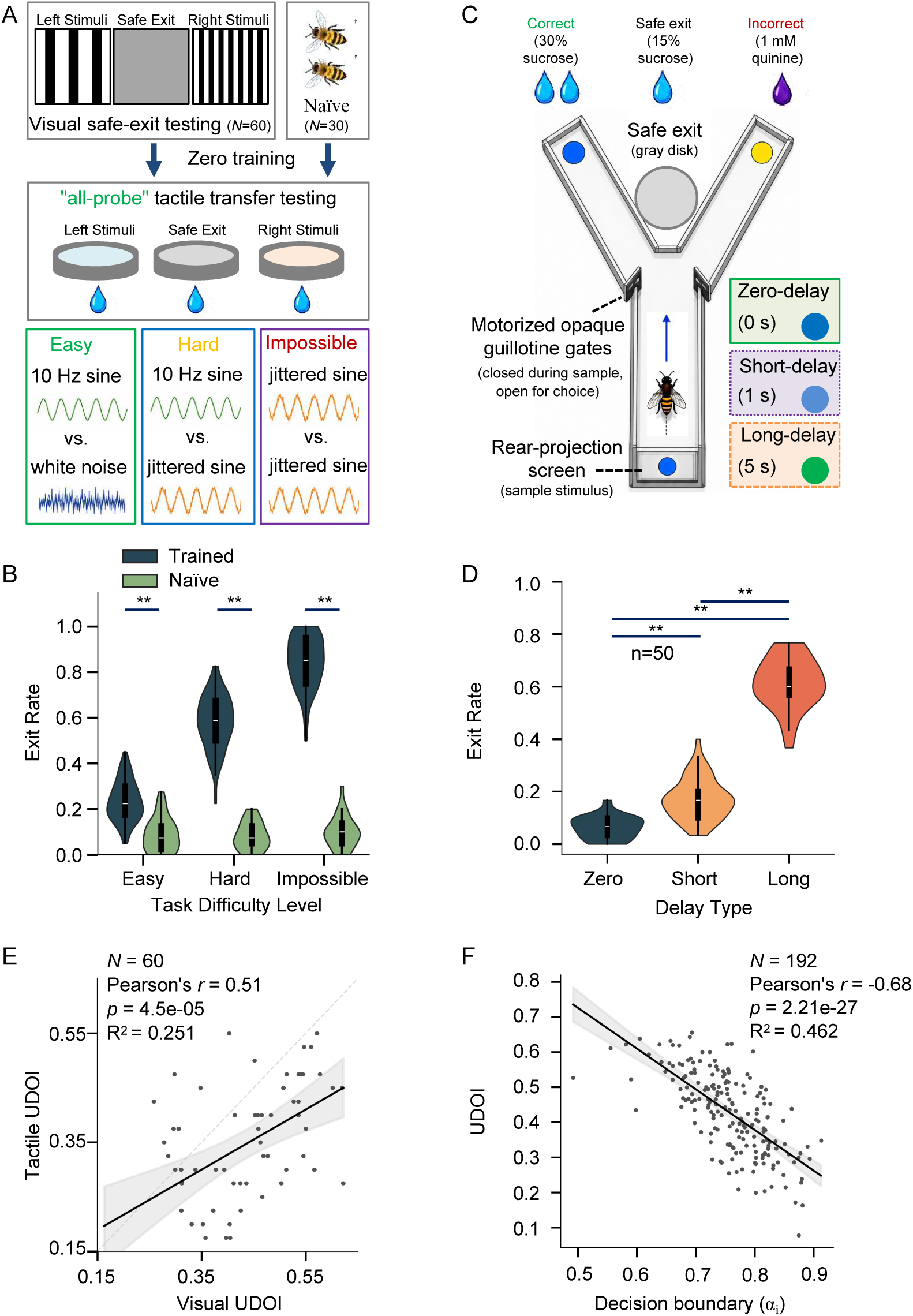
The opt-out strategy is an abstract, domain-general behavioral policy. (A) Cross-modal tactile transfer setup. Visual gratings were replaced by two vibrating platforms delivering tactile stimuli paralleling visual difficulty: Easy (10 Hz sine vs. noise), Hard (sine vs. jittered sine), Impossible (identical jittered waves). Bees received no tactile training; all choices yielded the same small reward (“all-probe” design). (B) Immediate transfer to touch. Visually trained bees (*n* = 60) showed a clear difficulty-dependent gradient: Easy (23.5 ± 8.7%), Hard (58.1 ± 11.6%), Impossible (84.4 ± 11.3%) (GLMM, Hard vs. Easy: *Z* = 21.8, *p* < 0.001). Naïve bees (*n* = 30) showed uniformly low, difficulty-insensitive exit rates (∼9.1 ± 5.1%). (C) Delayed match-to-sample (DMTS) task schematic. Bees viewed a sample color, endured a delay, then chose the matching color from two alternatives. (D) Transfer to memory uncertainty. Opt-out rates: Zero-delay (6.4 ± 4.3%), Short delay (17.1 ± 7.8%), Long delay (61.3 ± 9.5%) (GLMM, Long vs. Short: *Z* = 11.4, *p* < 0.001; Short vs. Zero: *Z* = 8.1, *p* < 0.001). (E) Uncertainty-Driven Opt-Out Index (UDOI; Hard-Easy opt-out difference) correlated across visual and tactile tasks (Pearson’s *r* = 0.51, *p* < 0.001), indicating stable individual traits. (F) Decision bound height α_i_ negatively correlated with UDOI. Pearson’s *r* = −0.68, *p* < 0.001, R^2^ = 0.46, *n* = 192. Bees with stricter thresholds (larger α_i_) opted out more on Easy trials, yielding lower UDOI.

Transfer was immediate and strongly difficulty-dependent: opt-out rates rose from Easy (23.5 ± 8.7%) to Hard (58.1 ± 11.6) to Impossible trials (84.4 ± 11.3%; GLMM, Hard vs. Easy: *Z* = 21.8, *p* < 0.001; Impossible vs. Hard: *Z* = 10.7, *p* < 0.001; Fig. 2B, Fig. S4A and B, and Table S7). In contrast, naïve bees (*N* = 30) showed uniformly low and difficulty-insensitive exit rates of approximately 9.1 ± 5.1% (Fig. 2B; Fig. S4 C and D). Thus, the transferred behavior depended on prior acquisition of the opt-out strategy, not on an innate response to tactile difficulty.

We next tested whether the same strategy extended from perceptual to mnemonic uncertainty. A new cohort (*N* = 50) was trained on a delayed match-to-sample (DMTS) task (Fig. 2C). To dissociate the opt-out strategy from “choose the middle” heuristic, the safe exit was positioned centrally ahead of the choice arms. Bees viewed a sample color, endured a delay, then chose the matching color from two alternatives. Delay length manipulated uncertainty: Short (1 s), Long (5 s, where memory decay increased uncertainty), and Zero-delay (sample visible throughout). After task acquisition, the safe exit was introduced. Exit rates were low on Zero-delay (6.4 ± 4.3%) and Short-delay trials (17.1 ± 7.8%; GLMM, *Z* = 8.1, *p* < 0.001), but increased sharply on Long-delay trials (61.3 ± 9.5%; GLMM, Short vs. Long: *Z* = 11.4, *p* < 0.001; Fig. 2D, Fig. S4E, and Table S8). This pattern demonstrates that the opt-out strategy generalized from perceptual ambiguity to uncertainty arising from internal memory degradation.

### Individual differences in uncertainty monitoring are stable across modalities

If the opt-out strategy reflects a domain-general form of uncertainty monitoring, then individual sensitivity should show correlate across tasks. To test this, we computed an Uncertainty-Driven Opt-Out Index (UDOI) (*11*) for each bee as the difference in opt-out rate between Hard and Easy trials. Individual UDOI values in the visual task were positively correlated with those in the tactile task (Pearson’s *r* = 0.51, *p* < 0.001; Fig. 2E and Table S9), suggesting that uncertainty-monitoring propensity is a stable individual trait rather than a transient response to a specific sensory context.

To formalize the decision process, we developed a multialternative race model with three parallel accumulators (target, non-target, safe exit), incorporating expected value computation and an optional confidence threshold (*9, 23, 24*). The full model, which integrates both economic value and uncertainty sensitivity, outperformed five alternatives (pure difficulty, reinforcement learning, reaction time, stimulus similarity, and value-based without threshold; ΔWAIC > 50, Bayes Factor > 1000; see Methods and Table S10). Hierarchical Bayesian fitting (Stan) (*25, 26*) confirmed successful convergence for all 192 bees (R^ < 1.01; see Methods and Fig. S5A,B). Uncertainty sensitivity was primarily determined by the confidence threshold height *α_i_* (range 0.49-0.91, mean = 0.76), which strongly negatively correlated with UDOI (Pearson’s *r* = −0.68, *p* < 0.001, R^2^ = 0.46; Fig. 2F). Non-decision time *t_0i_* showed minimal variation (CV = 4%; Fig. S5C,D), and drift rates *υ_i_* clustered near the lower bound (*υ*_min_ = 0.05; Fig. S5E), indicating negligible contributions from sensorimotor speed. Thus, the observed behavior can be explained by an online estimate of evidence quality modulated by individual ambiguity tolerance, beyond mere economic optimization.

### Liquid neural network captures temporal dynamics of uncertainty-guided choices

To examine how uncertainty manifests in the temporal structure of behavior, we trained a Liquid Neural Network (LNN) (*27*), a continuous-time recurrent network, to predict opt-out choices from behavioral time series. Input features excluded the externally provided task difficulty label to prevent information leakage. The model achieved an ROC-AUC of 0.787 (Fig. S6A) and an average precision of 0.827 on held-out trials (Fig. S6B), indicating that the temporal structure of behavior contains a robust decodable signature of decision uncertainty. Control analyses confirmed that this performance was not driven by decision latency alone: a latency-only logistic regression yielded AUC = 0.65, and the LNN retained significant predictive power when trained on fixed-length sequences or after removing latency information (see Methods). Learned time constants (τ) clustered tightly around 2.93 across three independently trained models (uncertainty monitoring, information seeking from scratch, and information seeking via transfer learning), approaching the dynamic upper limit of 4.0 (Fig. S6C–E). This preference for long-term integration was invariant across task difficulty levels (Fig. S6F–H), suggesting that behavioral policy relies on sustained temporal integration of evidence rather than a rapid difficulty-triggered response.

We next tested generalization across tasks. Initializing an information-seeking LNN with weights from the uncertainty monitoring model improved recall (0.646 vs. 0.543, Δ = +10.3%) and F1-score (0.654 vs. 0.589) compared with training from scratch (Fig. S6I), indicating shared temporal features between opt-out and information-seeking paradigms. In zero-shot cross-modal transfer, the frozen visual uncertainty monitoring LNN reproduced the difficulty-dependent gradient in tactile data from trained bees (Fig. S6J) but not from naïve bees (Fig. S6K), indicating that the network learned strategy-specific temporal signatures rather than generic stimulus features. The same frozen model also generalized to the DMTS working memory task (Fig. S6L). The LNN is best understood as a computational description of the decision policy, rather than a mechanistic model of its neural substrate. Together, these analyses provide strong evidence that uncertainty-guided choice in bumble bees depends on a learned, transferable and domain-general decision policy.

## DISCUSSION

### From cognitive content to cognitive monitoring

Landmark studies on social signal learning, affective contagion, abstract rhythm perception, and spontaneous problem-solving have established that insect brains can represent, compute, and share rich cognitive content (*4–7*). Our study answers a fundamentally different question: whether bees can adjust their behavior based on the reliability of that content (*28*). We provide evidence that bumble bees possess an abstract, domain-general uncertainty-monitoring strategy, shifting the inquiry from what insect brains can represent to whether they can monitor the reliability of those representations (*29*).

### A stringent all-probe test of transferable uncertainty monitoring

The all-probe design is the methodological strength of this study. During tactile transfer, all choices, including both vibration platforms and the safe exit, yielded identical small rewards. Bees therefore could not learn which tactile stimulus was correct or more valuable within the test session. Difficulty-dependent opt-out behavior had to arise from a strategy acquired prior to the tactile test, rather than new reinforcement within it. This design separates transfer of a decision policy from new task learning and offers a generalizable framework for probing uncertainty-guided control in species where verbal reports or extensive training are not possible (*17,28*).

### A learned and transferable computational strategy

A drift-diffusion model incorporating a confidence threshold captured accuracy, reaction times, and graded opt-out rates across all difficulty levels, suggesting that bees track the quality of their own evidence accumulation during decision formation. The observed *αᵢ* – UDOI correlation is consistent with a causal role of the confidence threshold in driving metacognitive sensitivity, although causal proof would require neural manipulation. Liquid neural network analyses further showed that opt-out decisions could be predicted from temporal behavioral dynamics, that longer-timescale integration characterized the behavioral policy, and that transfer succeeded in trained bees but failed in naïve ones. This failure indicates that the network captured the temporal signature of an acquired uncertainty-monitoring strategy, not a generic response to stimulus difficulty. While these analyses support the learned and domain-general nature of the strategy, the temporal signatures are most parsimoniously interpreted as reflecting the behavioral expression of metacognitive monitoring, though we cannot fully exclude a broader evidence-accumulation process.

### Evolutionary and neural implications

Our results reveal that a compact insect neural architecture can support behavior consistent with uncertainty-guided decision-making (*30,31*). This extends the potential evolutionary origins of uncertainty monitoring, a core computational component often linked to consciousness, back by over 600 million years (*32,33*), to the last common ancestor of insects and vertebrates (*8,12,13*). Rather than implying a single evolutionary origin, it shows that uncertainty monitoring can emerge from small nervous systems and may represent a general computational solution for decisions under uncertain evidence (*22*). Mushroom bodies and the central complex represent plausible targets for future work on how uncertainty signals are generated and deployed (*34–36*).

### What the evidence does and does not show

This study provides behavioral and computational evidence for uncertainty-guided control in an invertebrate (*21*). Our claim is functional (*32*) rather than phenomenological. Bumble bees regulated behavior according to uncertainty, but these experiments do not establish whether they subjectively experience uncertainty. This work does not address subjective consciousness in bees. Rather, it provides strong evidence that behavior consistent with uncertainty monitoring can be generated by a brain with fewer than one million neurons and no cortex (*24, 25*).

By demonstrating behavior consistent with uncertainty monitoring in a brain of remarkable simplicity (*37*), our work suggests that the minimal neural substrate required for uncertainty-guided decision-making may be far smaller than previously assumed. This finding suggests that a core computational capacity often linked to higher-order cognition can emerge from compact neural architectures— an insight that invites a fundamental reconsideration of the evolutionary biological foundations of cognitive monitoring (*14,38*).

## ACKNOWLEDGMENTS

**Funding:** This work was supported by the grants from the Guangdong Basic and Applied Basic Research Foundation (2024A1515030060 to L.Y.), Shenzhen Key Industry R&D Program (202509013000454 to L.Y.), Chinese National Natural Science Foundation Projects (81602422 to L.Y.). **Author contributions:** Conceptualization: L.Y.; Methodology: Y.H., L.Y.; Investigation: Y.H., Q.Y., Q.W., F.Y., S.W.; Visualization: L.L., J.X., C.W.; Funding acquisition: L.Y.; Supervision: F.Y., L.Y.; Writing – original draft: Y.H., L.Y.; Writing – review & editing: Q.Y., Q.W., F.Y., S.W., L.Y. **Competing interests:** The authors declare that they have no competing interests.

## Supplementary Materials

### Materials and Methods

#### Subjects and Husbandry

A total of 470 commercially obtained, colony-raised worker bumble bees (*Bombus terrestris*) from 12 queen-right colonies (Biobest, Belgium) were used, matching the established model system in foundational work (*4, 39*). All sample sizes across every experimental cohort were determined a priori based on effect sizes derived from pilot experiments (Cohen’s d > 0.8 for Hard vs Easy opt-out) to achieve statistical power > 0.95 at α = 0.05. The final *N* = 192 for the visual uncertainty monitoring task (including the information-seeking sub-task), *N* = 60 for the punishment removal control, *N* = 60 for the cross-modal tactile transfer (trained subset), *N* = 30 for the naïve tactile control, and *N* = 50 for the DMTS memory task—all independently met or exceeded the required sample size benchmarks calculated from the same power analysis framework. No post-hoc sample size adjustments were made. Colonies were housed in standard wooden nest boxes (28 × 16 × 10 cm) connected to a dedicated flight arena (60 × 40 × 30 cm) and maintained under controlled environmental conditions (28 ± 2°C, 50-60% relative humidity) on a reversed 12:12 h light:dark cycle. During the dark (subjective day) period, colonies and experimental rooms were illuminated by red LED light strips (peak λ= 660 nm), to which bees are relatively insensitive, thereby minimizing uncontrolled visual cues during experimentation and colony maintenance. Bees were provided with *ad libitum* natural pollen (approx. 3g deposited in the nest box daily) and 15% (w/w) sucrose solution via gravity feeders in the flight arena. This diet was maintained throughout the study period, except during specific testing hours. Prior to experiments, individual bees were briefly cold-anaesthetized on a Peltier cooling plate and uniquely marked on the thorax with colored, numbered Opalithplättchen tags. Only active, non-naïve foragers—bees observed voluntarily and repeatedly collecting sucrose from feeders in the common arena for at least 48 hours prior to testing—were selected for behavioral testing. This husbandry and selection protocol ensures a consistent, motivated, and healthy subject pool, a critical prerequisite for complex cognitive testing. All procedures were approved by the Institutional Human Research Ethics and Animal Care and Use Committee at Peking University and Peking University Shenzhen Hospital.

### General Apparatus: Visual Discrimination Flight Tunnel

#### Design and Rationale

The core behavioral apparatus was a linear flight tunnel (100 cm L × 25 cm W × 20 cm H; internal dimensions) constructed from 10mm-thick black PVC panels, chosen for its rigidity and light-blocking properties. The internal walls and ceiling were lined with matte gray paper (60% reflectance) to create a uniform, non-reflective visual field, isolating the experimental stimuli. This controlled environment is essential for studying visual discrimination and decision-making without confounding cues. One end featured a computer-controlled, opaque sliding gate for controlled bee entry. The opposite end housed the decision zone: a vertical panel containing three circular, transparent acrylic landing platforms (3 cm diameter, 0.5 cm thick) arranged in a linear equidistant arrangement (6 cm, center-to-center). This spatial configuration presents a clear, simultaneous three-alternative choice.

#### Stimulus Presentation and Reward Delivery

Each platform was backlit by an independently controlled, diffused LED panel (Adafruit NeoPixel grids, 60 LEDs/dm^2^) for precise visual stimulus presentation, allowing instantaneous changes. Reward and punishment were delivered via a custom-built, computer-controlled 4-channel fluidics system using solenoid valves (The Lee Company) and calibrated microcapillary tubes. Sucrose (see concentrations below) and quinine (1 mM quinine hydrochloride) solutions were dispensed in precise volumes (5 µL) onto the center of a platform within 100 ms of a bee’s sustained (>0.5 sec) contact, ensuring tight temporal contiguity between choice and consequence—a cornerstone of effective operant conditioning.

#### Data Acquisition

A high-resolution, high-speed camera (Basler acA2040-90um) mounted 50 cm directly above the decision zone, recorded all trials at 90 fps. The entire apparatus was housed inside a light-tight, sound-attenuating cabinet lined with acoustic foam to exclude external visual and auditory disturbances. This integrated design ensures precise experimental control and high-fidelity behavioral recording. Illumination for video recording was provided by two infrared LED arrays (850 nm) mounted on the ceiling.

#### Visual Stimuli

Stimuli were vertical, square-wave black-and-white gratings (100% Michelson contrast) generated in PsychoPy. The discriminated parameter was spatial frequency. The “Target” stimulus was a grating of 0.1 cycles/mm. To manipulate decision difficulty and thus internal uncertainty, the “Non-target” stimulus varied: “Easy” trials used a highly discriminable 0.3 cycles/mm grating; “Hard” trials used a poorly discriminable 0.12 cycles/mm grating; “Impossible” trials presented two identical 0.1 cycles/mm gratings, making perceptual discrimination impossible. The “Safe Exit” option was a uniform gray disk of matched mean luminance. This carefully calibrated difficulty gradient is a standard and necessary feature for interpreting behavioral opt-out as a function of uncertainty (*10, 13*).

### Uncertainty Monitoring with a Passive “Safe Exit”

#### Feasibility & Rationale

This paradigm operationalizes “uncertainty-guided decision-making” by offering a guaranteed, low-value alternative to a risky, high-value discrimination. Its feasibility is grounded in bees’ proficient visual learning and their ability to associate distinct spatial locations with outcomes. The design’s power lies in testing whether an insect’s choice between these options is dynamically guided by an internal, trial-by-trial estimate of evidence strength or uncertainty, rather than a fixed preference.

#### Pre-training & Apparatus Familiarization

Individual marked bees were allowed to freely forage in the flight tunnel for 30 minutes with all three platforms presenting the uniform gray disk, each dispensing 5 µL of 15% sucrose upon landing. This established the apparatus as a reward source and neutralized any initial side biases.

#### Two-Choice Discrimination Training

Bees were then trained on the forced-choice visual discrimination (Target vs. Non-target) without the exit option (the central platform location was covered by a blank, matte gray insert to prevent its association as a choice option at this stage). A session consisted of 100 trials. A correct choice (landing on Target) was rewarded with 5 µL of 30% sucrose. An incorrect choice was punished with 5 µL of 1 mM quinine solution, a potent but harmless aversive stimulus for bees. “Easy” and “Hard” trials were randomly interleaved (50 each per session). Training continued until a bee achieved a criterion of >80% correct on Easy trials over a rolling window of 50 consecutive Easy trials. Performance on Hard trials during the final training session was 52.1% ± 3.2% (mean ± SD), confirming it was near chance level. This ensured not only task learning but also direct experience of the difficulty gradient, a prerequisite for forming an uncertainty signal.

#### Forced-Choice Control Experiment (Impossible Trials without Safe Exit)

To distinguish uncertainty-guided decision-making from a lower-level response-conflict account, we conducted a forced-choice control. If opt-out on Impossible trials merely reflects an irresolvable choice conflict, then when the safe exit is unavailable, bees should exhibit substantially prolonged decision latencies, frequent mid-flight returns, and chance-level accuracy on Impossible trials. Conversely, if bees monitor uncertainty and can still commit to a random choice when forced, their latencies on Impossible trials should be comparable to those on Hard trials (where evidence is weak but not absent), and accuracy should remain at chance.

#### Subjects

A new cohort of 60 trained bumble bees was used. Sample size was determined a priori (power > 0.95 at α = 0.05, Cohen’s d > 0.8 for latency difference between Hard and Impossible, based on pilot data).

#### Test phase

Immediately after reaching criterion, each bee completed a single session of 90 forced-choice trials (30 Easy, 30 Hard, 30 Impossible, pseudorandomly interleaved with no more than four consecutive trials of the same difficulty). On every trial, the bee flew from the start gate to the two-choice array. Landing on the target (correct) was rewarded with 5 μL of 30% sucrose; landing on the non-target (incorrect) was punished with 5 μL of 1 mM quinine. No safe-exit option was ever present. The inter-trial interval was 30 s.

#### Data recording

For each trial we recorded: (i) Decision latency (ms): time from crossing the start-gate line to sustained contact (>500 ms) with either platform. (ii) Choice accuracy: correct (target) or incorrect (non-target). (iii) Mid-flight return: a binary score (1 if the bee left the start gate, flew into the decision zone, and then returned to the start gate without landing on any platform within 10 s; 0 otherwise). Trials with a return were excluded from latency and accuracy analyses but counted separately. All scoring was performed offline by two independent observers blind to trial difficulty (inter-rater reliability > 99%). This forced-choice control established that, in the absence of any exit option, bees treat Impossible trials as random (accuracy ∼50%, latencies comparable to Hard trials), thereby ruling out a pure response-conflict explanation and providing a clean baseline against which the subsequent safe-exit opt-out behavior can be interpreted as uncertainty-driven.

#### Introduction of the Safe Exit Option

Following stabilization of discrimination performance (for the main cohort, *N* = 192), bees entered a brief “Safe Exit Introduction” phase. The central platform was uncovered and permanently displayed the uniform gray disk. In non-test shaping phase ( ≤ 10 trials per bee), the standard punishment for incorrect discriminations was temporarily suspended. landing on the new gray platform (Safe Exit) was rewarded with 5 µL of 15% sucrose solution, while correct/incorrect choices on the main task continued to be rewarded/punished as in training. This allowed bees to discover the new option’s value and consequence without the confounding pressure of punishment.

#### Formal Test Phase

Full task contingencies were reinstated. Each trial now presented a simultaneous three-alternative choice. Choosing the Safe Exit delivered 5 µL of 15% sucrose and immediately ended the trial. Test sessions (100 trials per bee) consisted of randomly interleaved Easy, Hard, and Impossible trials (40, 40, and 20 trials, respectively). Bees typically completed one full session per day. The specific reward concentrations were chosen based on prior preference tests in our lab confirming that bumble bees reliably prefer 30% > 20% > 15% sucrose. In a separate two-choice preference test (*N* = 20 bees), bees significantly preferred 30% over 20% sucrose solution (choice proportion 82.5 ± 5.8%; binomial test, *p* < 0.001), and also preferred 20% over 15% sucrose (choice proportion 78.3 ± 6.2%; binomial test, *p* < 0.001). During the test phase, we additionally quantified antennal investigation. The number of times the bee’s antennal tip contacted the stimulus panel surface was counted within the decision-latency window (from stimulus onset to final choice). The Antennal_Tapping_Frequency was calculated as the total number of contacts divided by the decision latency in seconds, normalizing investigation effort to decision duration.

### Punishment Removal Control Experiment

#### Subjects

A separate cohort of visually trained bumble bees (*N* = 60) was used. All bees were naïve to the punishment-removal manipulation and were individually marked for identification. Housing and maintenance conditions were identical to those described in the main experiment.

#### Apparatus and Stimuli

The same flight tunnel, visual stimuli and two-choice discrimination apparatus were used as described in the main opt-out paradigm (see main text). The safe exit option was always available and consisted of a separate landing platform leading to a guaranteed reward (5 µL 15% sucrose in both phases; see below).

#### Procedure

The experiment comprised two consecutive phases performed on the same set of bees in a single session (approximately 3 hours). Between phases, bees were allowed a 5-min rest period in a holding chamber with access to water. No retraining or re-familiarization was provided between phases.

Phase I-Baseline (with punishment). Bees first completed 90 trials (30 per difficulty level: Easy, Hard, Impossible) using the standard opt-out paradigm. In each trial, the bee chose either to discriminate between the two gratings or to take the safe exit. If the bee discriminated and chose the target (correct), it received 5 µL 30% sucrose; if it chose the non-target (incorrect), it received 5 µL of 1 mM quinine solution. The safe exit always delivered 5 µL 15% sucrose. Trials were pseudorandomly interleaved with equal probability per difficulty. Trials were pseudorandomly interleaved with the constraint that no more than five trials of the same difficulty occurred consecutively. The inter-trial interval was 30 s, during which the bee returned to the start gate.

Phase II-Punishment removal. Immediately following Phase I, the same bees underwent another 90 trials (again 30 per difficulty) in which all punishment was eliminated. On every trial: (i) if the bee discriminated (chose either target or non-target), it received 5 µL 15% sucrose regardless of correctness; (ii) if the bee took the safe exit, it also received 5 µL 15% sucrose. Thus, the reward magnitude for all outcomes was identical (5 µL 15% sucrose), removing any differential cost for errors. The physical layout, stimulus presentation, and trial timing were identical to Phase I.

#### Data Recording

A choice was scored when the bee made sustained contact (>500 ms) with a landing platform. An ‘opt-out’ was defined as a sustained landing on the central gray platform (safe exit). Choices were scored offline by two independent observers blind to the experimental condition using custom Python scripts; inter-rater reliability exceeded 99%. For each trial we recorded: subject ID, trial number, difficulty, phase (Baseline or NoPunishment), choice (discriminate or opt-out), and if discriminated, whether the chosen stimulus was correct (target) or incorrect (non-target). Opt-out rates were calculated per bee per difficulty per phase.

#### Statistical Analysis

All statistical analyses were performed in R (v4.3.1) using the lme4 package (v1.1.34) for GLMM fitting and the emmeans package (v1.8.9) for post-hoc comparisons. We fitted a generalized linear mixed model (GLMM) with binomial error distribution and logit link to the binary opt-out response. Fixed effects included difficulty (Easy, Hard, Impossible), phase (Baseline, NoPunishment), and their interaction. Bee ID was included as a random intercept to account for repeated measures.. Post-hoc pairwise comparisons were performed using Tukey’s HSD with a significance threshold of α = 0.05. All reported p-values are two-sided.

### Active Information Seeking at a Cost

#### Feasibility & Rationale

This paradigm probes a more cognitively demanding, active form of uncertainty monitoring control—the capacity to recognize insufficient information and take deliberate action to resolve it. Moving beyond passive uncertainty avoidance, it requires the animal to perform a deliberate, sequenced action that incurs a tangible cost to resolve uncertainty. This provides stronger evidence against low-level explanations, such as a generalized aversion to difficult stimuli or simple escape responses. The design is feasible because bumble bees are proficient at learning multi-step sequences and forming associations between a transient cue and a delayed reward.

#### Apparatus Modification

The linear flight tunnel was partitioned into two sequential decision zones by a transparent, vertical acrylic gate that could be lowered remotely. Zone 1 contained a single landing platform designated as the “Information Request” platform. Zone 2, located 20 cm further along the tunnel, contained the standard two-choice array (Target vs. Non-target platforms). The apparatus ensured that bees would naturally traverse Zone 1 before reaching Zone 2. The gate was initially open.

#### Sequence Training (Establishing the Behavioral Chain)

Before introducing any cost, bees underwent a crucial Sequence Training phase to establish the core “information request” action and its consequence. At this stage, bees were already proficient in the visual discrimination task in Zone 2 but had no experience with Zone 1. Guided Introduction: Bees were allowed to fly freely in the modified tunnel. Landing on the Zone 1 platform triggered an unambiguous, 500-ms predictive cue: the LED panels beneath platforms in Zone 2 flashed at 5 Hz, highlighting the one displaying the correct Target stimulus for that trial. Chain Establishment: After the cue, the bee then would proceed to Zone 2 and make its choice. A correct post-cue choice (matching the cued stimulus) was rewarded with the full, high-value reward (5 µL of 30% sucrose). An incorrect choice was punished with quinine. Objective & Outcome: This training had two key objectives: (a) To forge a strong cause-and-effect link between the voluntary act of landing on the Zone 1 platform and the subsequent resolution of uncertainty in Zone 2. (b) To instill a stable behavioral chain: “Land on Zone 1 platform → Trigger informative cue → Fly to Zone 2 → Use cue to choose correctly → Receive high reward.” This chain became the baseline, trained response against which the cost manipulation in the test phase could be evaluated.

#### Test Phase with Economic Cost

Following sequence training, the critical test phase introduced an economic trade-off to assess strategic information-seeking. The contingency for landing on the Zone 1 “Information Request” platform was changed: it now yielded no immediate reward but still triggered the cue. If a bee requested information (landed in Zone 1) and then made a correct choice in Zone 2, it received a discounted reward (5 µL of 15% sucrose). An incorrect choice in Zone 2 was still punished. Crucially, bees could also “skip” the information request by flying directly from the entrance to Zone 2. A correct choice made without requesting information was rewarded with the higher, undiscounted reward (5 µL of 30% sucrose). An incorrect choice was punished. Thus, “requesting information” now carried a guaranteed opportunity cost (forfeiting the higher reward), modeling a real-world decision to invest resources in acquiring knowledge.

#### Critical Control: Random Free-Cue Trials

To rule out non-cognitive explanations—such as a simple attraction to the flashing cue or a habitual preference for the Zone 1 platform unrelated to information need—a critical control condition was interleaved in 20% of the test trials. In these “Random Free-Cue” trials, the predictive cue was automatically presented as soon as the bee’s body centroid crossed into the front half of Zone 1, independent of whether it landed on the platform. All other task contingencies remained identical: the bee still had to make a choice in Zone 2 for a reward or punishment.

Logic of the Control: This design creates a diagnostic dissociation. If bees land on the Zone 1 platform merely because they are attracted to the cue itself, they should land less often on Free-Cue trials (since the cue is already provided) compared to regular trials. Conversely, if landing is a strategic, need-based action, its pattern should change specifically: landing should be strongly suppressed on Impossible trials where the cue is worthless, but may persist on Easy/Hard trials due to residual habit from training, even though the cue is free. This control directly tests the intentionality and judgment underlying the information-seeking act.

#### Testing Protocol

Each bee completed 100 test trials. Trials (Easy, Hard, and Impossible difficulties in Zone 2) and types (80 regular cost trials and 20 Random Free-Cue trials) were randomly interleaved. We recorded the bee’s choice in Zone 1 (to seek information or not) and its subsequent discrimination accuracy in Zone 2.

### Cross-Modal Generalization of the “Uncertainty” Rule (Vision to Touch)

#### Feasibility & Rationale

This is the most decisive test for an abstract, domain-general uncertainty-monitoring rule. Bees possess sophisticated tactile vibration sensitivity. Testing the immediate transfer of the “opt-out when uncertain” strategy to a novel, untrained sensory modality with an “all-probe” design (equal reward for all choices) is a stringent and feasible test. It prevents new associative learning during the test, forcing the animal to rely solely on a pre-existing, abstract behavioral policy.

#### Tactile Apparatus

The LED panels in the decision zone were replaced with two custom vibration platforms. Each consisted of a 3 cm diameter acrylic landing surface glued to the cone of a 2” loudspeaker (Visaton FRWS 5-8 Ohm). Speakers were driven by a stereo amplifier (Topping PA3s) connected to a digital function generator (Rigol DG1022Z). Vibration displacement was measured with a laser vibrometer (Polytec OFV-5000) and calibrated to a peak-to-peak amplitude of 50 ± 5 µm for all stimuli.

#### Tactile Stimuli

“Target” vibration: a 10 Hz pure sine wave. “Non-target (Easy)”: 0-100 Hz band-limited white noise. “Non-target (Hard)”: a 10 Hz sine wave with a 180° phase reset (“jitter”) occurring at random intervals (mean: 50ms, range: 30-70 ms), creating perceptual similarity. “Impossible“: two identical jittered sine waves. To confirm that the tactile Easy condition was indeed perceptually discriminable, we conducted a pilot discrimination test with a separate group of naïve bees (*N* = 30) under full-reward feedback. These bees achieved 86.7 ± 8.2% accuracy on Easy trials, confirming that the Easy tactile stimuli were readily discriminable. Hard trials yielded 52.3±9.1% accuracy, and Impossible trials yielded 49.2±7.8%.

#### Procedure

Bees (*N* = 60) that had completed the visual Safe Exit test (Exp. 1) were used. They received zero training or exposure to the tactile stimuli or their associations. In a single 100-trial test session, the Safe Exit option (gray disk) was present. The critical manipulation was that every choice outcome—landing on either vibration platform or the Safe Exit—delivered the same 5 µL of 15% sucrose solution. This eliminated any possibility of learning the tactile discrimination or its outcomes within the session. A separate, completely naïve control group of bees (*N* = 30, with no prior experience in any opt-out task) underwent the identical tactile test to establish a baseline, difficulty-insensitive response pattern.

### Cross-Domain Generalization to a Memory-Based Task

#### Feasibility & Rationale

To assess the generality of the uncertainty-monitoring strategy, we tested whether it transfers to uncertainty arising from an internal cognitive source (memory decay) rather than perceptual ambiguity. A delayed match-to-sample (DMTS) task is a well-established paradigm for studying working memory in bees and introduces a quantifiable source of internal uncertainty (the fading memory trace over time).

#### DMTS Apparatus

A modified Y-maze (stem: 20 cm long; each arm: 15 cm long, 8 cm wide, 10 cm high) was constructed from matte white acrylic. The stem contained a rear-projection screen for presenting the sample stimulus (a blue or yellow circle, 4 cm diameter). At the end of each arm was a landing platform. Motorized, opaque guillotine gates (controlled by servo motors) separated the stem from the choice arms. A third, central platform serving as the “Safe Exit” was located 4 cm in front of the choice arm bifurcation.

#### Initial Shaping

A new cohort of bees (*N* = 50), with no prior experience in the visual or tactile uncertainty monitoring tasks, was trained to fly into the stem, where the sample stimulus was presented. Upon presentation, the gates to both arms opened. Both choice platforms initially displayed the matching color. Landing on either was rewarded (5 µL, 30% sucrose). This established the basic “go to matching color” rule.

#### Discrimination Training

Once bees reliably entered the stem, the choice was made differential. One arm platform now displayed the matching color, the other the non-matching color. A correct match choice was rewarded; an incorrect non-match choice was punished with quinine. The sample color and correct side were randomized. Training continued until a bee achieved >75% correct over 40 consecutive trials.

#### Introduction of Delay

A delay interval between sample offset and gate opening was introduced, starting at 0.5s and gradually increasing to 1s over several sessions. Performance was stabilized at >70% correct at the 1s delay before proceeding.

#### Introduction of the Safe Exit

The central Safe Exit platform (gray disk) was introduced with a short, punishment-suspended shaping phase (∼10 trials), where choosing the Safe Exit yielded the low reward (5 µL, 15% sucrose) to familiarize bees with the new option.

#### Formal Test Phase

In the critical test, delay intervals were varied pseudo-randomly: Short Delay (1s, “Easy”), Long Delay (5s, “Hard”), and Zero-delay Control (sample remained on until choice, making a memory error impossible). Choosing the Safe Exit always delivered the low reward (5 µL, 15% sucrose). A correct match after a delay delivered the high reward (5 µL, 30% sucrose). Each bee completed 90 test trials (30 per condition). The key dependent measure was the opt-out rate as a function of delay length, with the prediction of significantly higher opt-outs in Long Delay compared to Short Delay and Zero-delay trials.

### Computational Modeling: Drift-Diffusion Model (DDM) with a Confidence Threshold

#### Rationale

To move beyond descriptive statistics and provide a mechanistic, quantitative account of the choice data (proportions and reaction times for correct, error, and opt-out responses across all difficulty levels), we developed a computational model that directly mirrors the tri-choice structure of the experimental task.

#### Model Architecture

Instead of a two-stage process (first choose A vs. B, then decide to opt-out), we implemented a parallel three-accumulator race model that simultaneously competes for a response: one accumulator for choosing the target (A), one for the non-target (B), and a third accumulator for the safe exit option. Each accumulator integrates evidence over time until one reaches its respective decision threshold. The drift rates for the A and B accumulators depend on the stimulus similarity (higher similarity → lower drift rate), while the safe exit accumulator integrates a constant positive value reflecting the guaranteed small reward (15% sucrose). Critically, we incorporated an expected value computation into the decision rule: the decision to commit to a choice (target, non-target, or safe exit) is made when the accumulated evidence for that option exceeds a threshold, and the final choice is determined by whichever accumulator hits its threshold first. The expected value of committing to the target or non-target is computed as *EV* = *P*(correct)×*R*_correct_−*P*(error)×*R*_error_, where *P*(correct) is estimated from the current state of evidence (using the relative height of the winning accumulator), *R*_correct_ =30% sucrose, and *R*_error_ = quinine punishment(*40*). The safe exit has a fixed *EV* equal to the guaranteed 15% sucrose. The model thus naturally produces opt-out behavior when the expected value of gambling is lower than the safe option, without requiring an explicit uncertainty-based confidence threshold. However, to test the uncertainty monitoring hypothesis, we also consider a variant where a confidence threshold (a separate parameter) can trigger an earlier opt-out even when the *EV* of the gamble is still positive, reflecting a non-economic sensitivity to uncertainty. The height of the confidence threshold is a fitted parameter capturing individual willingness to tolerate ambiguity.

#### Model Fitting and Comparison

All models were fit using hierarchical Bayesian estimation implemented in Stan (*26*). Parameters were estimated with weakly informative priors (e.g., Normal (0, 1) for drift rates on log scale; Gamma (2, 0.5) for thresholds). Individual-level parameters were drawn from group-level normal distributions with estimated means and standard deviations, accounting for inter-bee variability. Model fitting was performed using Hamiltonian Monte Carlo with 4 chains, 2000 warm-up iterations, and 4000 sampling iterations per chain. Convergence was verified via R^ <1.01. Model comparison was conducted using Watanabe-Akaike Information Criterion (WAIC) and leave-one-out cross-validation (LOO-CV) via the loo package in R. To evaluate the necessity of the uncertainty-sensitive component, we compared our full model (with both EV and confidence threshold) against a battery of alternative models: (i) Pure Difficulty Model: Opt-out rate depends solely on external task difficulty (Easy/Hard/Impossible), with no internal evidence accumulation. (ii) Reinforcement Learning Model: Bees learn an opt-out policy based on past rewards using a simple Q-learning algorithm, without any uncertainty tracking. (iii) Reaction Time Model: Opt-out is triggered when decision latency exceeds a fitted threshold, irrespective of stimulus or confidence. (iv) Stimulus Similarity Model: Opt-out probability is a direct function of the physical similarity between the two stripe patterns. (v) Value-Based Tri-Choice Model (without confidence threshold): Uses only expected value maximization with no additional uncertainty-related threshold. (vi) Full Model (with confidence threshold): Includes both expected value and a confidence threshold. Each model’s WAIC and LOO scores were computed; the model with the lowest score (best predictive accuracy) was considered the preferred explanation. Additionally, we computed Bayes Factors using the bridge sampling method (*41*)to compare the full model against the best alternative.

#### Computational Modeling of Decision Dynamics

To move beyond associative statistical descriptions and probe the temporal dynamics and potential computational mechanisms underlying the observed uncertainty-guided behaviors, we employed a Liquid Neural Network (LNN) model. LNNs are a class of continuous-time recurrent neural networks particularly suited for modeling sequential decision processes that evolve based on an internal state. In contrast to Generalized Linear Mixed Models (GLMMs), which quantify static associations and statistical significance between variables, the LNN aims to simulate and generate the observed behavioral sequences, allowing us to infer the properties of the latent dynamical system that could produce such behavior. The primary goals of this analysis were twofold: (i) to test whether the behavioral time series contained a robust, machine-learnable signal related to decision uncertainty, independent of external difficulty labels, and (ii) to characterize the computational properties—such as its effective timescale—of the simulated decision system.

#### Data Preprocessing and Sequence Construction

Raw behavioral trial data from the uncertainty monitoring test (Table S2_Three-choice decision-making trials) and the active information-seeking experiment (Table S6_Active information seeking trials) were used. For each trial performed by an individual bee, we constructed a multi-feature time-series vector. Crucially, to avoid information leakage, the feature vector did not include the externally provided task difficulty label. Instead, the features were limited to those that could be observed by an external observer or inferred from the bee’s own behavior: Stimulus identity (one-hot encoding of target side), Trial type (’Regular’ vs. ‘Random_Free_Cue’ trials), Historical action (one-hot encoding of bee’s action in the previous timestep), and Cumulative reward (normalized trace of recent reward history). The length of each sequence was defined by the trial’s actual decision latency. The terminal label for each sequence was the bee’s final choice for that trial: opt-out (safe exit) vs. continue (choose A or B). In total, approximately 19,200 valid behavioral sequences were constructed.

#### Control for Latency Confound

Because decision latency is known to correlate with difficulty and opt-out probability, we implemented three control analyses to isolate the contribution of temporal dynamics beyond mere reaction time: (i) Latency-only Logistic Regression: A logistic regression model using only the trial’s decision latency as a predictor of opt-out. (ii) Fixed-Length Sequences: All sequences were padded/truncated to a uniform length (the median latency across all trials), removing the possibility that the model simply uses sequence length as a proxy for latency. (iii) Latency-Removed LNN: The LNN was trained on sequences where the latency information was removed (i.e., sequences were not ordered by real time but instead resampled at a fixed number of steps per trial, and the absolute time duration was not provided).

#### Dataset Partitioning

To ensure rigorous evaluation and simulate generalization to novel individuals, we performed a stratified split by bee ID, creating three mutually exclusive subsets: **Training Set**: 70% of bees’ data (∼13,440 sequences), used to update model parameters; **Validation Set**: 15% of bees’ data (∼2,880 sequences), used for hyperparameter tuning, monitoring overfitting, and implementing early stopping; **Test Set**: 15% of bees’ data (∼2,880 sequences), serving as a held-out set of “novel bees” that never influenced training. All final reported performance metrics (e.g., AUC) are derived from this independent test set.

#### Liquid Neural Network Architecture and Training

### Model Structure

We employed a single LNN layer with 32 hidden units. The dynamics of the hidden state h(t) are governed by the ordinary differential equation: dh/dt = (1/T)⊙[-h + *f* (Wx + Uh + b)], where ***f*** is the hyperbolic tangent (tanh) activation function, **T** is a vector of learnable time constants governing dynamical speed, ⊙denotes element-wise multiplication, and **W**, **U**, and **b** are input weights, recurrent weights, and bias, respectively. The hidden state is read out by a linear layer followed by a sigmoid to produce a probability for the binary decision (opt-out/continue). **Key Parameter-Time Constant (τ)**: A central feature of our model is that the time constant τ was a learnable, neuron-specific parameter. This allows the network to adaptively discover dynamical patterns operating at different timescales to best explain the behavioral data. **Training Details**: The model was trained using the Adam optimizer with an initial learning rate of 0.001 and a cosine annealing schedule. The loss function was binary cross-entropy. Training was performed with a batch size of 64. A strict early-stopping policy (patience=10) was applied, halting training when the validation loss failed to improve for 10 consecutive epochs and reverting to the best model checkpoint. Training was completed on a single NVIDIA RTX 4090 GPU in approximately 30 minutes.

### Evaluation Protocol and Metrics

Model performance was comprehensively evaluated on the independent test set using the following complementary metrics: **Area Under the Receiver Operating Characteristic Curve (ROC-AUC)**: Reported for opt-out prediction. An AUC > 0.5 indicates predictive power above chance, with values closer to 1.0 representing a stronger ability to decode an “uncertainty” signal from behavioral time series. **Precision-Recall Curve and Average Precision (AP)**: Given the imbalanced nature of opt-out vs. continue trials, the PR curve and AP were calculated to provide a more complete assessment of the model’s performance in identifying the positive class (opt-out). **Reliability Diagram and Expected Calibration Error**: The calibration plot compares predicted probabilities against actual observed frequencies. The ECE quantifies the average miscalibration, assessing the reliability of the model’s confidence in its predictions. **Confusion Matrix**: Standard performance metrics (accuracy, precision, recall) were calculated at the default threshold of 0.5 to analyze the model’s error profile. We report results for the primary LNN model (without difficulty label) and for the latency-control models.

### Ablation Study (Removing Task Difficulty Label)

To directly address the concern that the model might simply exploit the difficulty label, we performed an ablation study. We trained an LNN model with the difficulty label included (as in the original manuscript) and compared its performance to the model without the difficulty label. Furthermore, we evaluated the model’s ability to predict opt-out within each difficulty level separately (Easy, Hard, Impossible) using only behavioral features (no difficulty label). If the model can still discriminate opt-out vs. continue within a given difficulty level, it demonstrates that it extracts a genuine uncertainty signal beyond what is captured by the external difficulty.

### Model Interpretability and Mechanistic Analysis

To interpret the decision mechanism learned by the LNN, we conducted the following analyses: **Time Constant Distribution Analysis**: The learned time constants τ for all 32 hidden units were extracted and visualized. A τ < 1 indicates a fast dynamical unit reliant on recent inputs, while τ > 1 would indicate a slow unit capable of long-term integration. This distribution reveals the characteristic “internal clock” the model employed to explain bee behavior. **Behavioral Effect Recovery**: The trained model was probed with inputs corresponding to different task difficulties (Easy, Hard, Impossible). The model’s mean predicted opt-out rate for each condition was plotted, providing an independent verification of whether it recapitulated the fundamental task difficulty gradient observed in the bees. **Visualization of Individual Differences**: Based on the model’s processing of each bee’s behavioral sequences, bees were ranked and grouped (low, medium, high) by their mean predicted opt-out rate. This visualization captures the continuum of inter-individual variability in opting strategies. Additionally, we analyzed the **learned representations** to confirm that the model does not implicitly reconstruct the difficulty label.

### Rationale and Value of the LNN Analysis Beyond GLMM

The introduction of the LNN in this study is not intended to replace the GLMM but to provide a complementary, computational perspective. The GLMM has perfectly fulfilled its role by rigorously testing the statistical associations and significance between fixed effects (e.g., task difficulty, trial type) and the opt-out rate. However, the GLMM treats each trial as a relatively independent observation, making it difficult to fully capture the dynamic nature of decision-making as a continuous temporal process. In contrast, the LNN, by simulating a dynamical system capable of generating similar behavioral sequences, offers the following unique insights:

### Revealing Temporal Dynamics

By analyzing the learned time constants, the LNN suggests that the bees’ uncertainty assessment assessment may rely on a slow, long-timescale integration process (all τ > 1, clustered around 2.93) rather than rapid evidence decay. This provides a quantitative signature for the computational style of uncertainty monitoring.

### Quantifying Signal Strength

The AUC of 0.787 for opt-out prediction without difficulty label on the test set provides an objective, quantifiable piece of evidence indicating a high signal-to-noise ratio for decoding the “internal state of uncertainty” from behavioral time series. This strongly supports the view that behavior is a reliable indicator of internal cognitive states.

### Providing a Unifying Mechanistic Framework

The LNN architecture itself constitutes a potential computational mechanism hypothesis. We preliminarily demonstrated its extensibility to a multi-task model, using the same dynamical system to simultaneously simulate two manifestations of uncertainty-guided behavior—“opting out” and “information seeking”—laying the groundwork for constructing a more unified computational theory of uncertainty monitoring in the future.

### Offering New Analytical Dimensions

The LNN allows for the characterization of individual differences based on dynamic trajectories and enables analyses such as “counterfactual” simulations, which are difficult to achieve with traditional statistical models.

Therefore, incorporating LNN modeling into this manuscript significantly enhances the theoretical depth and methodological innovation of the work. It not only independently validates the core behavioral findings from a computational modeling perspective but, more importantly, elevates the discussion from “whether a behavior occurs” to “how the behavior might be generated through a dynamic computational process”. This builds a bridge connecting the uncertainty-guided behavior of bees with its underlying neural computational mechanisms.

### Data Analysis, Statistics and Availability

#### Behavioral Tracking

All video recordings were analyzed using DeepLabCut, a deep learning-based pose estimation toolkit, to extract precise coordinates for the bee’s body and head, as well as platform contact events. Custom Python scripts calculated decision latencies, flight paths, and investigation durations.

#### Statistical Modeling

Statistical analyses were performed using R v4.3.1 (lme4 v1.1-34, emmeans v1.8.9)), Python v3.10 (PyTorch v2.0.1, DeepLabCut v2.3.5), and Stan v2.32 (via the cmdstanr R interface), following and extending the rigorous analytical framework demonstrated in prior work (*4, 5*). Choice data (binary and multinomial) were analyzed using Generalized Linear Mixed Models (GLMMs) with a logit link function, incorporating individual Bee ID as random intercepts to account for repeated measures. Fixed effects varied by experiment but generally included trial difficulty, experimental group, and their interaction. For the cross-modal transfer experiment, the critical test was the Group (Trained vs. Naïve) × Difficulty interaction on opt-out rate. Reaction time data were analyzed using Gamma or Lognormal GLMMs with identical random-effects structures. For key GLMMs, model selection was guided by the Watanabe-Akaike Information Criterion (WAIC) and/or Leave-One-Out Cross-Validation (LOO-CV) (*42*). The final models reported were those with the lowest WAIC and highest expected log predictive density (ELPD) from LOO-CV. To examine whether uncertainty signals could be decoded from behavioral time series, we trained a Liquid Neural Network (LNN) using PyTorch. Model performance was evaluated on held-out trials using ROC-AUC and average precision (AP). Statistical significance of decoding performance was assessed via permutation tests (1,000 iterations) comparing the observed AUC against a null distribution obtained by shuffling trial labels. For the three primary GLMMs (visual opt-out, tactile opt-out, and DMTS opt-out), all pairwise post-hoc comparisons (Tukey HSD) were additionally evaluated using the Benjamini-Hochberg false discovery rate (FDR) procedure at *q* < 0.05. All comparisons reported as significant survived this correction. All reported *p*-values are two-sided, and confidence intervals are reported at the 95% level.

#### Reproducibility, Transparency and Availability

The study adheres to the MDAR (Materials Design Analysis Reporting) framework for reproducible science. All data supporting the findings of this study are available within the paper. All other data and materials supporting the findings of this study are available from the corresponding author on reasonable request. Source data are provided with this paper. The codes for statistical analysis are available at GitHub with the following link: https://github.com/Cuixiaojian21/bee_metacognition.

#### Ethical Statement

All procedures were conducted in accordance with the 3Rs principles (Replacement, Reduction, Refinement) and were approved by the Institutional Human Research Ethics and Animal Care and Use Committee at Peking University and Peking University Shenzhen Hospital. Bumble bees are not subject to specific animal welfare regulations in many jurisdictions, but all efforts were made to minimize potential distress. A brief cold anesthesia was used only for marking procedure. The sucrose and quinine solutions used are part of their natural foraging experience. Bees were housed in enriched colonies, provided ample food, and returned to their colonies after testing.

---

**Fig. S1.**
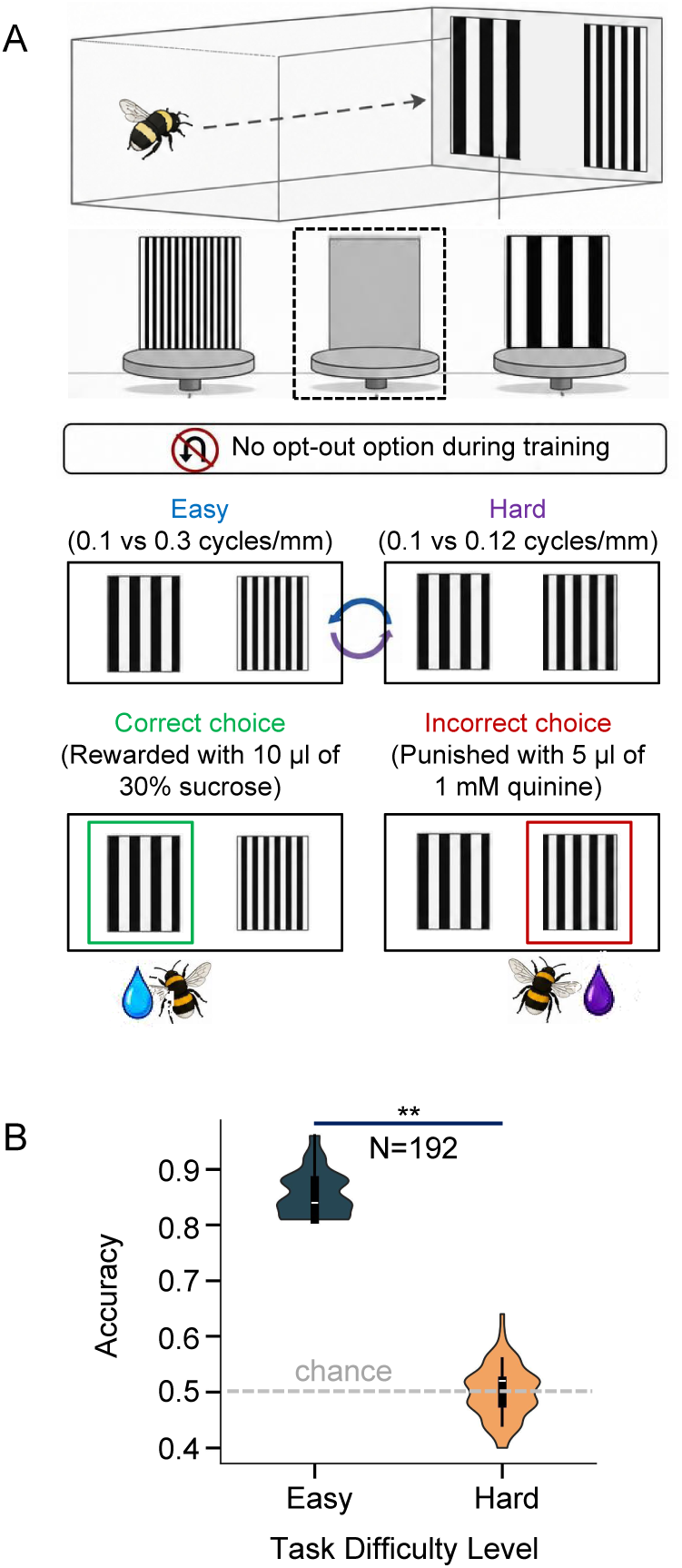
Training performance establishes a perceptual difficulty gradient. (A) Training phase schematic. Bees (*n* = 192) discriminated Target (0.1 cycles/mm) from variable Non-target gratings (Easy: 0.3 cycles/mm; Hard: 0.12 cycles/mm), randomly interleaved. Correct: 5 µL of 30% sucrose; incorrect: 5 µL of 1 mM quinine. No safe exit available. (B) Accuracy over final 50 training trials. Easy trials: mean = 0.856, median = 0.840 (above 0.5 chance). Hard trials: mean = 0.501, median = 0.520 (at chance). This confirms a robust difficulty gradient for probing uncertainty monitoring.

**Fig. S2.**
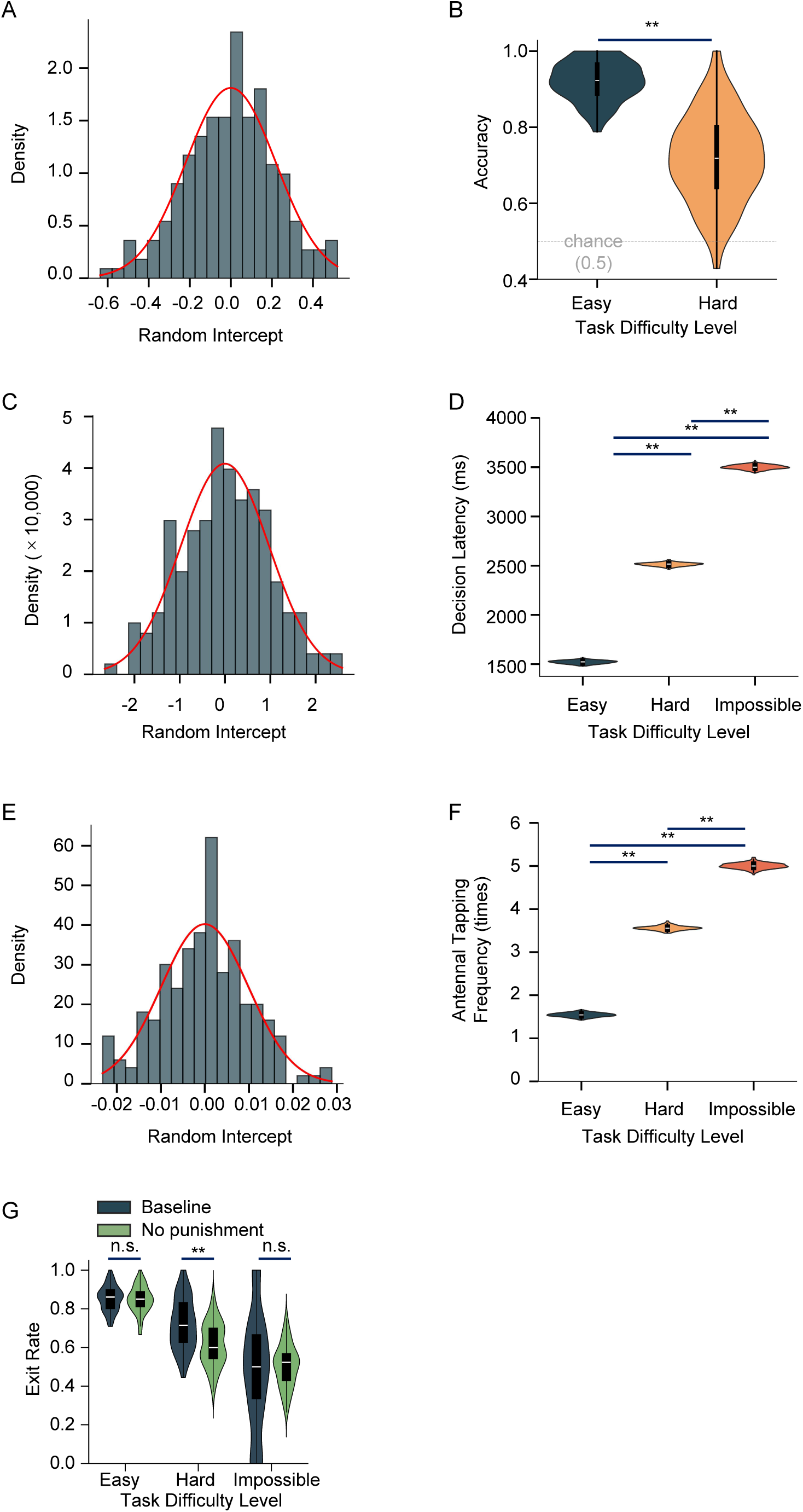
Behavioral indicators of uncertainty during decision-making (related to Fig.1A-C). (A) Random intercepts for opt-out behavior confirm stable individual differences (Mean = −0.0002, SD = 0.2209). (B) Accuracy on committed Hard trials (71.8±11.6%) significantly above chance (binomial test, *p* < 0.001), indicating confidence-based gambling. (C) Random intercepts distribution for the decision latencies in visual uncertainty monitoring task. (D) Decision latencies increased with difficulty: Easy (1518.8 ± 11.5 ms), Hard (2515.4 ± 11.5 ms) and Impossible (3514.9 ± 11.6 ms) consistent with extended evidence gathering under uncertainty. (E) Random intercepts distribution for the antennal investigations in visual uncertainty monitoring task. (F) Antennal investigation increased with difficulty: Easy (1.55 ± 0.34), Hard (3.56 ± 0.35) and Impossible (5.06 ± 0.45), indicating active perceptual assessment. (G) Accuracy during punishment removal control. Committed choice accuracy remained above chance on Hard trials (61.5 ± 12.4%; *p* < 0.001) but at chance on Impossible trials (50.3 ± 3.1%; *p* = 0.78). **, *p* < 0.001; n.s., not significant.

**Fig. S3.**
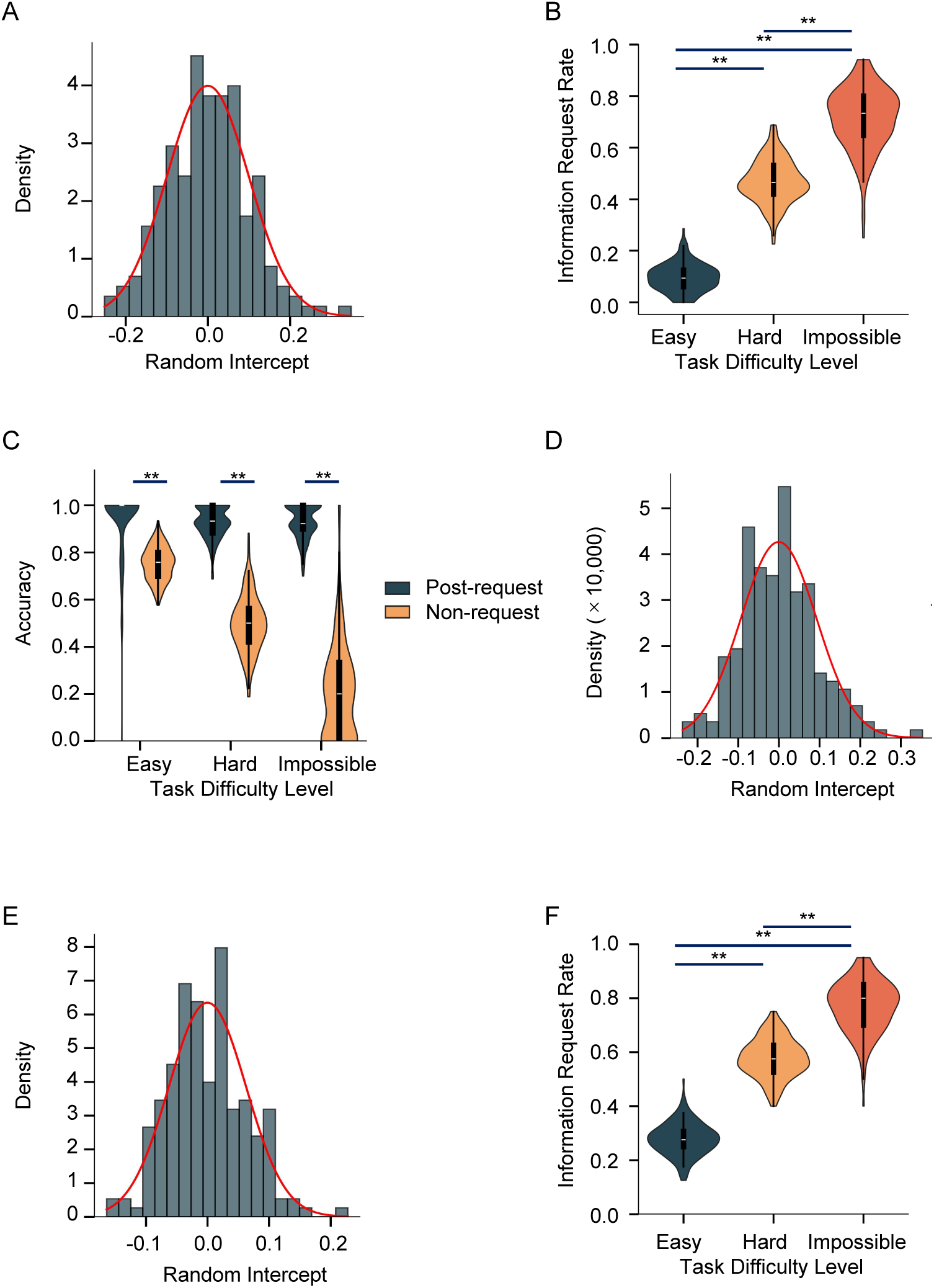
Strategic nature of active information seeking (related to Fig. 1D – F). (A) Random intercepts for information-seeking behavior in Regular (costly) trials. Distribution adhered to the theoretical normal curve (mean = −0.0001, SD = 0.1583), validating GLMM assumptions and confirming stable individual differences in information-seeking propensity. (B) Information request rates scaled with task difficulty in Regular trials. Requests increased stepwise from Easy (9.2 ± 3.0%) to Hard (45.6 ± 6.8%) to Impossible trials (70.8 ± 10.1%) (GLMM, all pairwise *p* < 0.001). (C) Information seeking effectively resolved uncertainty. After requesting information, discrimination accuracy exceeded 92% across all difficulty levels, transforming chance-level performance on Hard and Impossible trials into near-ceiling accuracy. (D) Random intercepts for information-seeking behavior in random free-cue trials. Distribution approximated the theoretical normal curve, confirming model validity. (E) Random intercepts for information-seeking behavior pooled across regular and random free-cue trials. Distribution conformed to the theoretical normal curve, supporting the robustness of the mixed-effects model. (F) Information request rates increased with difficulty when pooling both trials types. GLMM analysis (*Info_Requested∼Difficulty*+ (1|*Bee_ID)*) revealed a stepwise increase: Easy (27.7 ± 5.1%), Hard (57.8±5.6%), and Impossible (77.0±6.8%) (Tukey HSD, all pairwise *p* < 0.001).

**Fig. S4.**
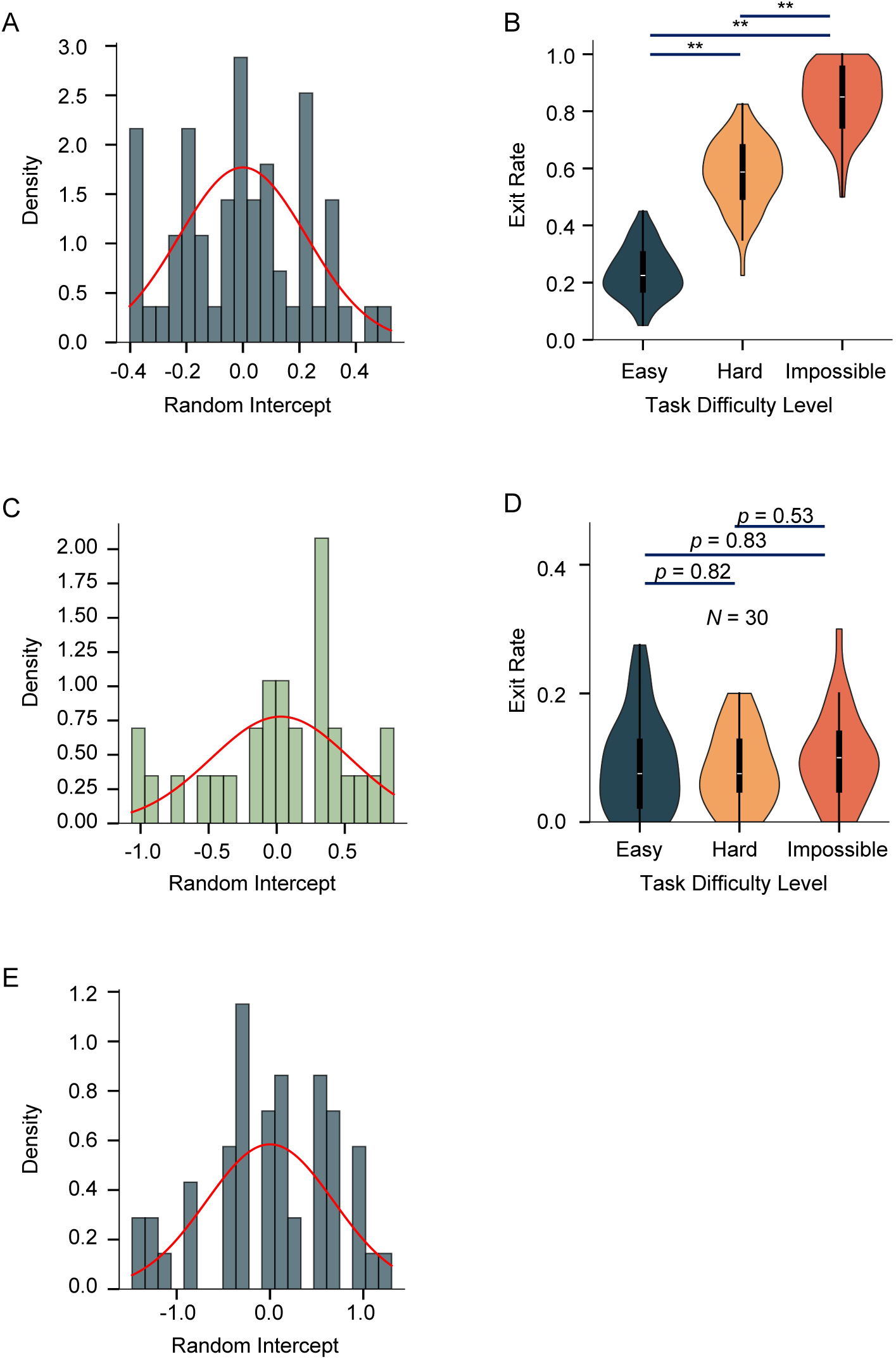
Cross-modal generalization confirms stable individual traits (related to Fig. 2A – D). (A) Random-effects validation for trained bees (*n* = 60) in the tactile test. Random intercept distribution (mean = 0.0002, SD = 0.2255) conformed closely to the theoretical normal curve, validating GLMM assumptions and confirming stable individual strategies. (B) Trained bees showed stepwise difficulty-dependent opt-out in the tactile task: Easy (23.5± 8.7%), Hard (58.1±11.6%), and Impossible (84.4±11.3%) (Tukey HSD, all pairwise *p* < 0.001), indicating zero-shot transfer of the uncertainty-monitoring rule to a novel modality without additional training or feedback. (C) Random-effects validation for naïve bees (*n* = 30) in the tactile task. Wider random intercept distribution (mean = 0.0298, SD = 0.5129) reflected greater behavioral volatility and the absence of a stable strategy. (D) Naïve bees exhibited uniformly low, difficulty-insensitive exit rates: Easy (9.0±1.1%), Hard (8.3±1.1%), and Impossible (9.8±1.4%) (Tukey HSD, all pairwise *p* >0.05). (E) Random-effects validation for the DMTS working-memory task. The near-zero random-effects variance indicated highly consistent strategies with negligible inter-individual differences. Distribution aligned with the normal curve, supporting robust model fit.

**Fig. S5.**
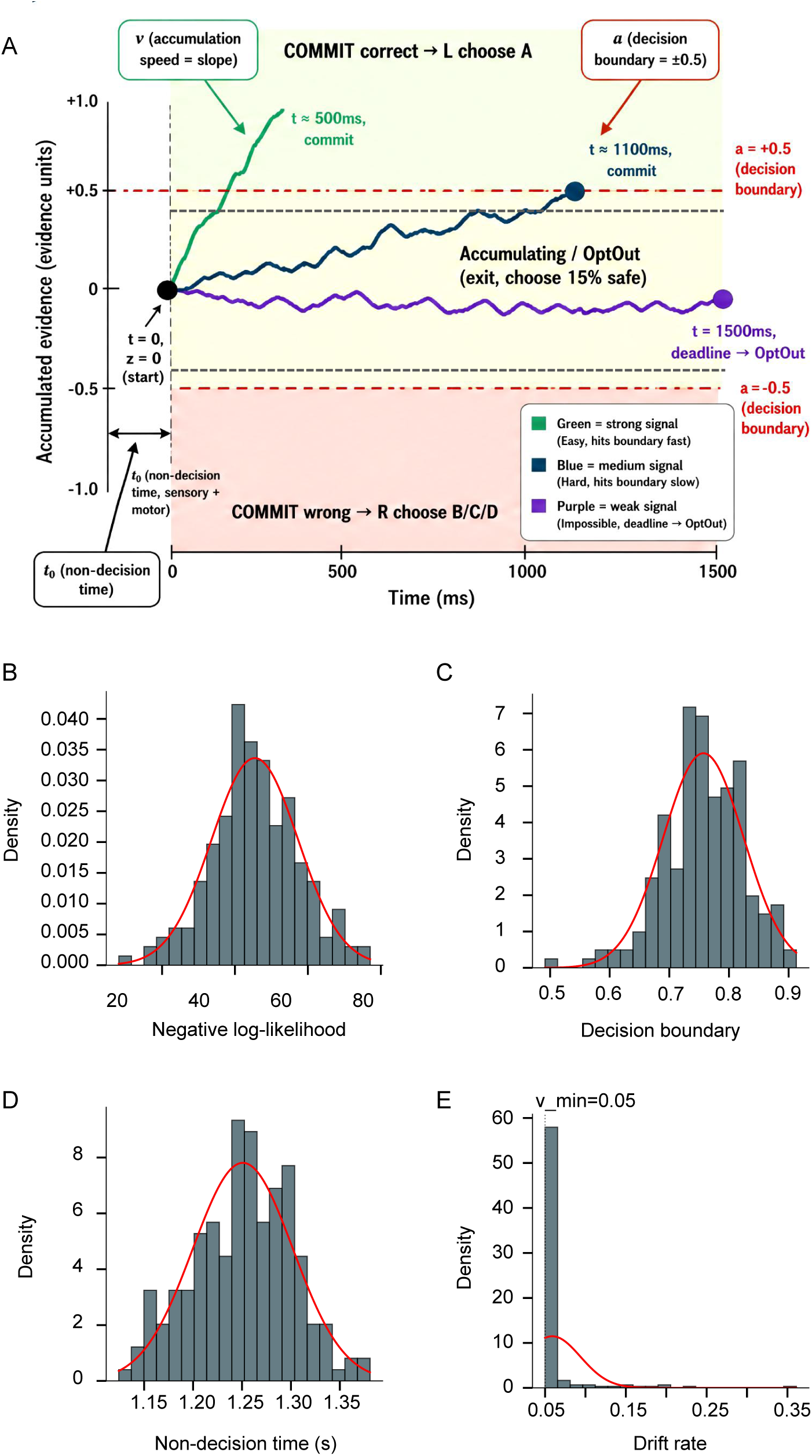
Drift-diffusion model (DDM) parameter distributions and relationship with uncertainty sensitivity. (A) Conceptual framework: Drift-Diffusion Model with confidence threshold. Evidence accumulation trajectories (colored lines) for Easy (blue), Hard (green), and Impossible (purple). Red dot-dashed lines: decision bounds (± *a*/2). Black dashed lines: confidence thresholds (± *c*). Easy trials cross the upper bound rapidly (high accuracy, low opt-out). Hard trials linger within the threshold zone (increased opt-out). Impossible trials hover near zero, triggering threshold and maximal opt-out. Shaded middle areas denote uncertainty zones. (B) Distribution of decision bound height (*α_i_*) across 192 bees. Range: 0.49-0.91, mean = 0.76. Higher *α_i_* indicates a stricter evidence threshold; within a fixed time window, insufficient accumulation drives opt-outs even on Easy trials. (C) Distribution of non-decision time (*t*_0*i*_). Range: 1.12-1.39 s, mean = 1.25 s, SD = 52 ms (CV = 4%). The minimal across-bee variation indicates similar sensorimotor processing; individual differences in uncertainty sensitivity arise primarily from *α_i_*, not *t*_0*i*_. (D) Distribution of drift rate (*υ_i_*). Most estimates clustered at the lower bound (*υ*_min_=0.05, gray dashed line), indicating minimal evidence accumulation within the fixed window. Low variance shows that individual differences stem from *α_i_* rather than sensory processing efficiency. (E) Distribution of negative log-likelihood (NLL) from per-bee DDM fits. NLL ranged from 8.2 to 77.1 with no convergence failures (R^< 1.01 for all 192 bees), confirming robust model fitting.

**Fig. S6.**
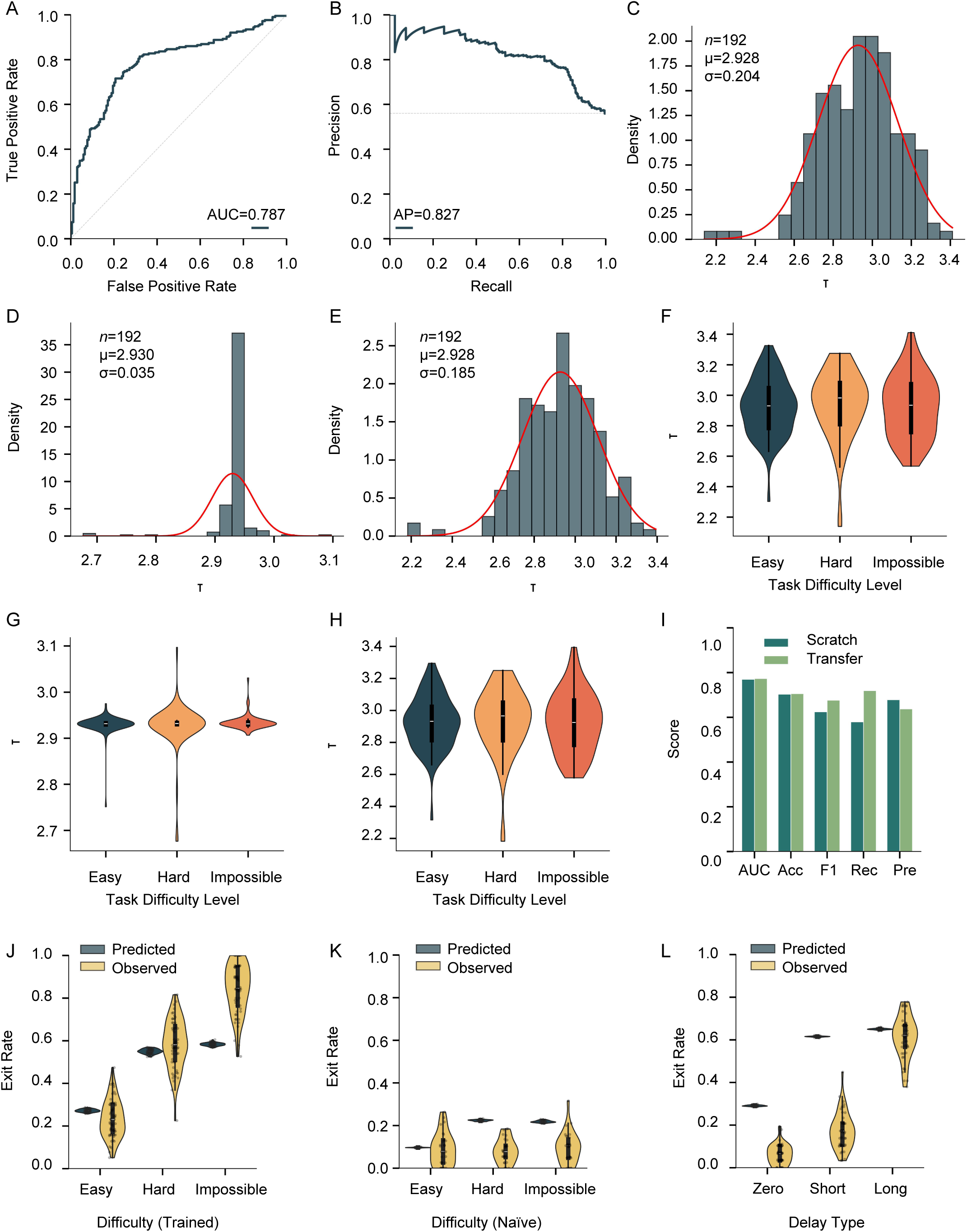
Liquid neural network reveals computational dynamics of uncertainty-guided decisions. (A) ROC curve for the Uncertainty monitoring model on the test set (*n* = 384, threshold = 0.5). AUC = 0.787. Steep rise at FPR < 0.2 indicates high true positive rates while controlling false positives. (B) Precision-recall curve for the same test set. AP = 0.827. Large area under the curve reflects favorable precision-recall tradeoff despite class imbalance. (C–E) Population distributions of learned τ at best epoch (histograms with Gaussian fits; n, mean *μ*, SD *σ* annotated). All three models converged to τ ∼2.93, approaching the dynamic upper limit (4.0), indicating long-timescale integration. The Uncertainty monitoring model showed the widest spread (C: *μ* = 2.928, *σ* = 0.204, τ ∼2.14-3.41); scratch Information Seeking the narrowest (D: *μ* = 2.930, *σ* = 0.035, τ ∼2.68-3.11); transfer Information Seeking intermediate (E: *μ* = 2.928, *σ* = 0.185, τ ∼2.22-3.38). (F–H) τ remains invariant across difficulty levels within each model (*n* = 192 per panel). Uncertainty monitoring (F) and transfer Information Seeking (H) preserved broad individual variation; scratch Information Seeking (G) collapsed toward the global mean, suggesting transfer retains individual-difference encoding. (I) Performance comparison of Information Seeking variants (Scratch vs. Transfer) on the test set (*n* = 384, threshold = 0.5). Transfer outperforms Scratch on all five metrics: AUC 0.777 vs. 0.770, Accuracy 0.708 vs. 0.677, F1 0.654 vs. 0.589, Recall 0.646 vs. 0.543, Precision 0.662 vs. 0.645. Largest gain in Recall (+10.3%); Precision increases modestly (+1.7%). (J–L) Zero-shot transfer of frozen Uncertainty monitoring model to external tasks. Predicted (dark blue) vs. observed (yellow) opt-out rates. (J) Tactile task (trained bees): both followed an “easy-low, hard-high” gradient (predicted: ∼0.27→0.58; observed: ∼0.23→0.84); close agreement for Easy and Hard trials (deviation ≤ 0.05). (K) Tactile task (naïve bees): observed flat near 0.10; predictions overestimated Hard (∼0.14) and Impossible (∼0.28), confirming the model captured a learned strategy, not generic difficulty responses. (L) DMTS task: predicted and observed followed a “short-low, long-high” gradient (predicted: ∼0.30→0.66; observed: ∼0.08→0.62).

## Notes

### Competing Interest Statement

The authors have declared no competing interest.

